# A breast Duct-on-Chip model for emulating invasive ductal carcinoma and testing therapies

**DOI:** 10.1101/2025.05.24.655087

**Authors:** Mohammad Jouybar, Jaap M.J. den Toonder

## Abstract

There are not many *in vitro* models that closely mimic ductal carcinoma *in vivo*. We present a novel breast duct-on-chip model designed to simulate ductal carcinoma *in situ* as well as invasive breast ductal carcinoma. This model features a tubular channel encapsulated in collagen and lined with normal epithelial cells. A basement membrane layer forms between the epithelium and the collagen, expressing basement membrane proteins. Invasive breast cancer cell lines are introduced into the breast duct to simulate ductal carcinoma. We observe two distinct lateral migration regimes that depend on the initial number of cancer cells within the duct. Additionally, the morphology of the breast duct and the invasion distance varies between the two regimes. Furthermore, we examine the effects of two widely used chemotherapies, Doxorubicin and Paclitaxel, on the invasion behavior of breast cancer cells within this model. The introduction of these drugs alters the invasion patterns of cancer cells, with Paclitaxel inhibiting invasion in 30% of tests. Notably, in the absence of treatment, all tests result in single-cell invasion. Our findings also highlight the dynamics of intraductal migration of cancer cells in non-invasive DCIS, revealing flow-like movements of cancer cells along the epithelial layer. This work establishes a valuable platform for investigating the mechanisms of cancer invasion from the epithelium and basement membrane, as well as evaluating chemotherapeutic strategies and testing novel cell-based therapies. It can have significant implications for understanding the dynamics of early-stage ductal carcinoma for improving treatment protocols of existing chemotherapies, and for developing and advancing novel therapies.

## Introduction

Cancer progression and fatal outcomes are rarely caused by the primary tumor, but by secondary tumors formed through metastasis. Cancer metastasis includes several consecutive steps: invasion, where cancer cells escape from the primary tumor through basement membranes and interstitial tissues; intravasation, in which cancer cells cross a blood vessel wall and enter the circulation; extravasation and seeding in distant organs; and tumor growth at the metastatic site [1,2]. The basement membrane (BM) is a thin sheet (ranging between ∼100 nm to 1.4 μm in thickness, depending on tissue type [3]), composed primarily of laminin and collagen IV, which plays a key role in normal tissue development and function [4–6]. The BM separates epithelial and endothelial cells from the connective tissue and is crucial for cell signaling, structural integrity, and barrier protection against cells and large molecules [5]. In the early stages of epithelial cancer (carcinoma), malignant cells proliferate within epithelial ducts but have not yet breached the epithelium or BM to invade surrounding tissue [5,7]. In breast cancer, this early stage is called ductal carcinoma *in situ* (DCIS) [8,9]. Once invasion occurs, cancer cells migrate into the surrounding stromal matrix, marking the transition to invasive ductal carcinoma (IDC) [8,9].

Numerous studies have examined breast cancer cell invasion both *in vivo* and *in vitro*; however, the early stages of metastasis, particularly cancer cell invasion from DCIS, are often overlooked. One reason is the lack of a suitable breast ductal model to study the dynamics of cancer cell invasion through the epithelium and BM. As far as we know, an accurate *in vitro* model of the BM does not currently exist. Previous studies have employed various hydrogels and techniques to mimic BM characteristics [10,11]. For instance, Wisdom et al. developed nanoporous, BM ligand-presenting interpenetrating network (IPN) hydrogels, enabling modulation of plasticity relevant to natural BM [10]. In our previous study, we developed a heterogeneous model for cancer invasion by encapsulating cancer cells in Matrigel droplets using a droplet-making system, creating an envelope resembling the BM. These droplets were sandwiched between collagen I layers to mimic the stromal matrix [11]. This model produced cancer invasion patterns different from those seen in single hydrogel conditions. The droplets were approximately 100 μm in diameter, resulting in a Matrigel layer thicker than physiological BM (100 nm to 1.4 μm) [3]. Current *in vitro* Organ-on-Chip (OoC) models also often fail to recreate an organized epithelial layer enveloping the breast cancer cells, as seen in DCIS *in vivo*. Nguyen et al. created parallel channels, one lined with endothelial cells as a blood vessel and the other containing cancer cells [12]. However, the epithelial layer was not included in their ductal channel. In another study, Kutys et al. developed a vascular and breast duct model comprising two tubular channels: one forming a perfused vessel and the other a dead-ended epithelial-lined duct positioned 500 μm away [13]. Their focus was on vessel–duct interaction, and they did not introduce cancer cells into the epithelial duct. Thus, a model that replicates the early stages of cancer metastasis is valuable for understanding disease progression and evaluating potential therapies.

One of the main goals of OoC systems has been to support drug testing while reducing animal testing. Several such models have already been used for this purpose [14–16]. Although OoC systems have advanced the recreation of organ microenvironments, disease modeling, and the study of cell-cell and matrix interactions, they are not yet fully leveraged for functional drug screening. We believe that establishing therapeutic testing methods at the laboratory level is essential to further integrate these systems into pharmaceutical workflows.

In this study, we developed a breast duct-on-chip model lined with breast epithelial cells and surrounded by a collagen matrix. A BM formed between the epithelium and collagen. Invasive breast cancer cell lines were introduced into the epithelial duct, allowing us to model DCIS on a chip for the first time. This innovative approach enabled us to emulate the transition from DCIS to IDC and to study cancer cell invasion through the epithelium and BM into the surrounding matrix. Furthermore, we used this model to evaluate the effects of chemotherapeutic agents Doxorubicin and Paclitaxel on morphology and invasion patterns of invasive breast ductal carcinoma within the chip. This study introduces a method for generating various ductal structures found in the human body and enables drug testing under diseased conditions.

## Results

### Development of a breast duct-on-chip model, expressing BM proteins

A breast duct-on-chip model was developed by forming a tubular channel within a collagen type I hydrogel (Figure 1a, b). This channel was seeded with normal breast epithelial cells (MCF10a), which adhered to the channel walls and formed a continuous epithelial lining, simulating a breast duct embedded within a 3D matrix (Figure 1b, c). The duct was connected to two reservoirs for cell seeding (Figure 1a (iii)) and two lateral reservoirs for media exchange, enabling nutrient diffusion through the hydrogel (Figure 1a (ii)). The chip was fabricated from polydimethylsiloxane (PDMS) using a stereolithography (SLA) 3D-printed mold [17,18], and bonded to a glass slide. Each reservoir was 5 mm in diameter and height, holding sufficient media to minimize evaporation during incubation. The duct channel had a diameter of 160 μm, created using a needle of the same size inserted into the chip before introducing the collagen gel. After collagen polymerization, the needle was carefully removed to form the hollow duct. MCF10a cells seeded into this channel first attached to the bottom and subsequently migrated to cover the entire lumen, forming a complete epithelial tube (Figure 1c, d; Supplementary video 1). Complete coverage of the channel was typically achieved within 17 hours using an initial seeding density of 2 × 10⁶ cells/mL (Figure 1c).

**Figure 1.**
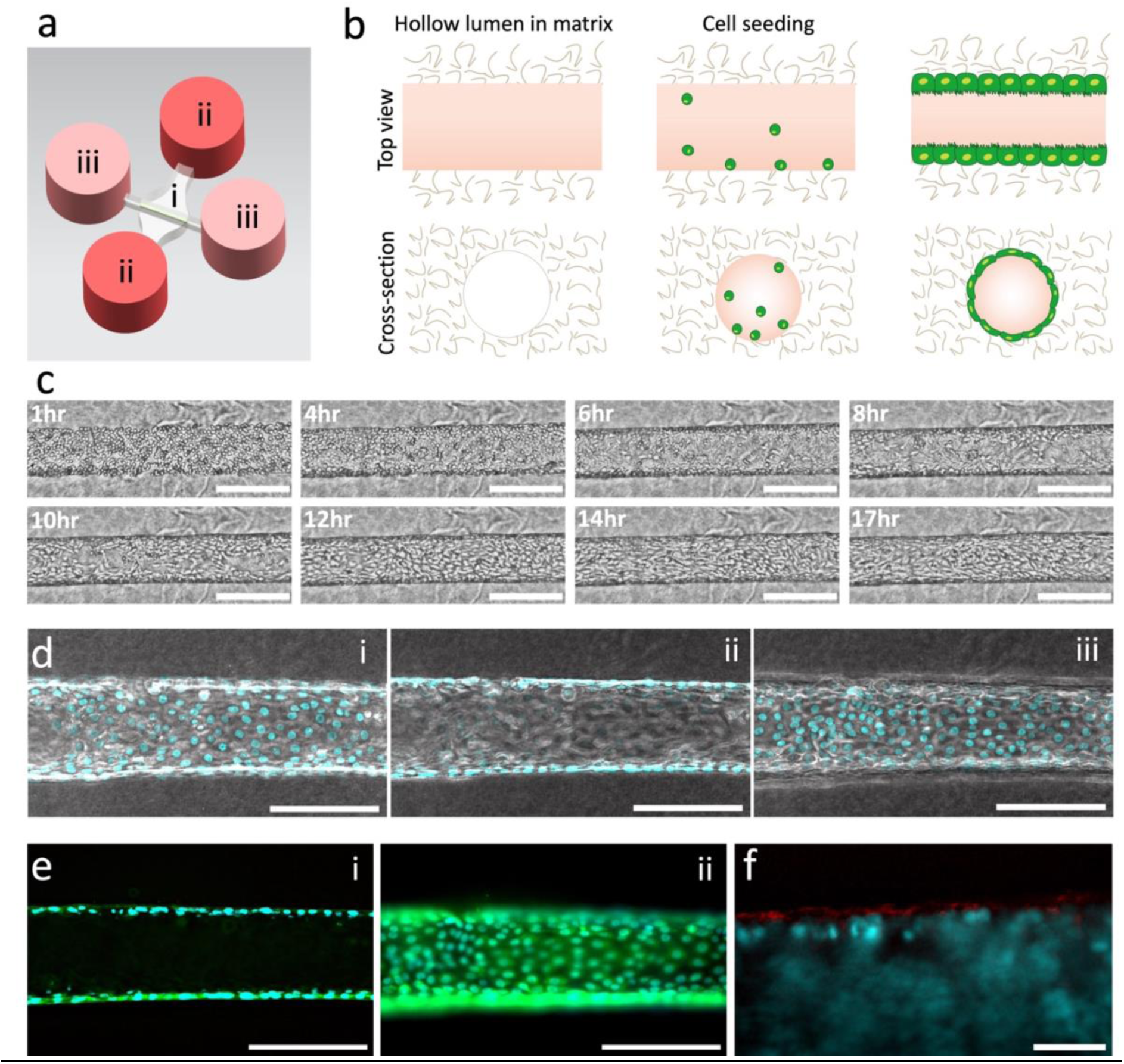
Development of a breast duct-on-chip model. a) Schematic of chip design. i = lumen, ii = reservoirs for gel infusion and media exchange (red), iii = reservoirs for cell seeding (pink), white = hydrogel. b) Schematic illustration of breast duct formation, from the hollow lumen in collagen I (left) to cell seeding (middle) and cell coverage (right). c) Phase contrast images of MCF10a cells in the breast duct from cell seeding to full coverage. Each image shows a snapshot of a time-lapse recording. The full coverage took 17 hours. Scale bars: 200 μm. See Supplementary video 1. d) Images of the (i) bottom, (ii) middle, and (iii) top of a breast duct-on-chip. The cell nuclei are shown in cyan. Scale bars: 200 μm. e) Fluorescent images of the (i) middle, and (ii) bottom of a breast duct-on-chip, stained for cell nuclei in cyan and laminin in green. Scale bars: 200 μm. f) Fluorescent image of the edge of a breast duct-on-chip in the interface with collagen gel, stained for cell nuclei in cyan and collagen IV in red. Scale bar: 30 μm.

Cancer metastasis involves the breaching of the epithelial layer and basement membrane (BM), allowing cancer cells to invade the surrounding stroma [5,7]. Several studies have aimed to replicate BM-like barriers in microfluidic models [19–27]. Porous membranes are commonly used in organ-on-chip (OoC) systems to mimic such barriers [19], but flat synthetic membranes often fail to capture the complexity of native BM architecture. PDMS membranes and electrospun hydrogel fibers have been widely explored due to their structural similarity to BM [20–25], and have been incorporated in epithelial-endothelial barrier models such as Lung-on-Chip and Gut-on-Chip [26,27]. However, the stiffness and thickness of synthetic PDMS membranes make them non-physiological [19]. Electrospinning offers potential for fabricating fibrous membranes with controlled porosity, but the elastic modulus of these materials often differs significantly from that of native BM [28]. Moreover, handling and integrating ultra-thin electrospun membranes (e.g., <10 μm) into microfluidic systems remains a technical challenge [19,23–25].

To address these limitations, biological hydrogels such as collagen I, fibrin, and Matrigel have been used within microfluidic devices to more closely replicate *in vivo* cell-matrix interactions [28–30]. These materials provide native ECM proteins and tunable mechanical properties. For example, Arik et al. demonstrated that collagen I can be dried to form a thin film that mimics flat BM membranes [31]. Matrigel, in particular, contains several native BM components [32,33], although recreating BM structure on curved or tubular geometries remains difficult. In our model, key BM proteins, including laminin (Figure 1e) and collagen IV (Figure 1f), were expressed at the interface between the epithelium and collagen gel. This was achieved by pre-coating the collagen-lined channel with Matrigel prior to epithelial cell seeding, which promoted cell adhesion and enhanced BM formation. Furthermore, incorporating the luminal geometry of the organ microenvironment into *in vitro* models is essential, as features like luminal channels are known to influence cellular morphology and orientation [34–38]. As a result, our duct-on-chip incorporates multiple *in vivo*-like features: a 3D tubular geometry, a continuous epithelial lining, and a self-assembled BM, bringing it a step closer to replicating the physiological breast duct environment.

### Modeling invasive breast ductal carcinoma on chip

We utilized the breast duct-on-chip system to investigate the early stages of breast cancer metastasis, focusing on the transition of cancer cells through the epithelial barrier and into the surrounding extracellular matrix. Conventional *in vitro* models, such as tumor spheroids embedded in hydrogels, typically involve a single cell type [39–42]. Even in multicellular contexts, these models often fail to replicate the formation of a well-organized epithelial layer surrounding the cancer cells [42]. Organoid-based models can form physiologically relevant epithelial boundaries with the matrix [39,43]; however, they do not allow controlled, sequential introduction of additional cell types, such as invasive cancer cells, after epithelial structure formation.

In our model, we introduced MDA-MB-231 breast cancer cells into the breast duct-on-chip system to study the transition from ductal carcinoma *in situ* (DCIS) to invasive ductal carcinoma (IDC). We compared this condition to a control in which MDA-MB-231 cells were seeded into a channel lacking an epithelial lining (Figure 2a). In the control setting, cancer cells invaded extensively into the surrounding collagen matrix (Figure 2b). In contrast, when seeded within an epithelialized duct, MDA-MB-231 cells disseminated into the matrix by breaching through the epithelial barrier (Figure 2c, d). Disruption of E-cadherin expression was observed in regions with a high concentration of cancer cells (Figure 2e, f), suggesting that epithelial integrity was compromised during invasion. Cancer cell invasion behavior was influenced by the initial cell seeding density. At low cancer cell densities, MDA-MB-231 cells formed clusters within the duct lumen and did not initially breach the epithelial barrier, mimicking early-stage DCIS (Figure 3a, Supplementary video 2). Over time, cancer cells proliferated within the duct (Figure 4a), contributing along with epithelial proliferation to a gradual increase in duct diameter (Figure 4b). Interestingly, this condition led to the emergence of multicellular epithelial protrusions extending from the duct into the collagen matrix, observed as early as day 2 (Figure 3a, 7.4c). These protrusions, although originating from regions adjacent to cancer cells, did not include MDA-MB-231 cells themselves. Cells at the invasion front displayed an elongated morphology, indicative of mesenchymal-like migration. This suggests that indirect signaling between cancer cells and neighboring epithelial cells may influence collective epithelial invasion.

**Figure 2.**
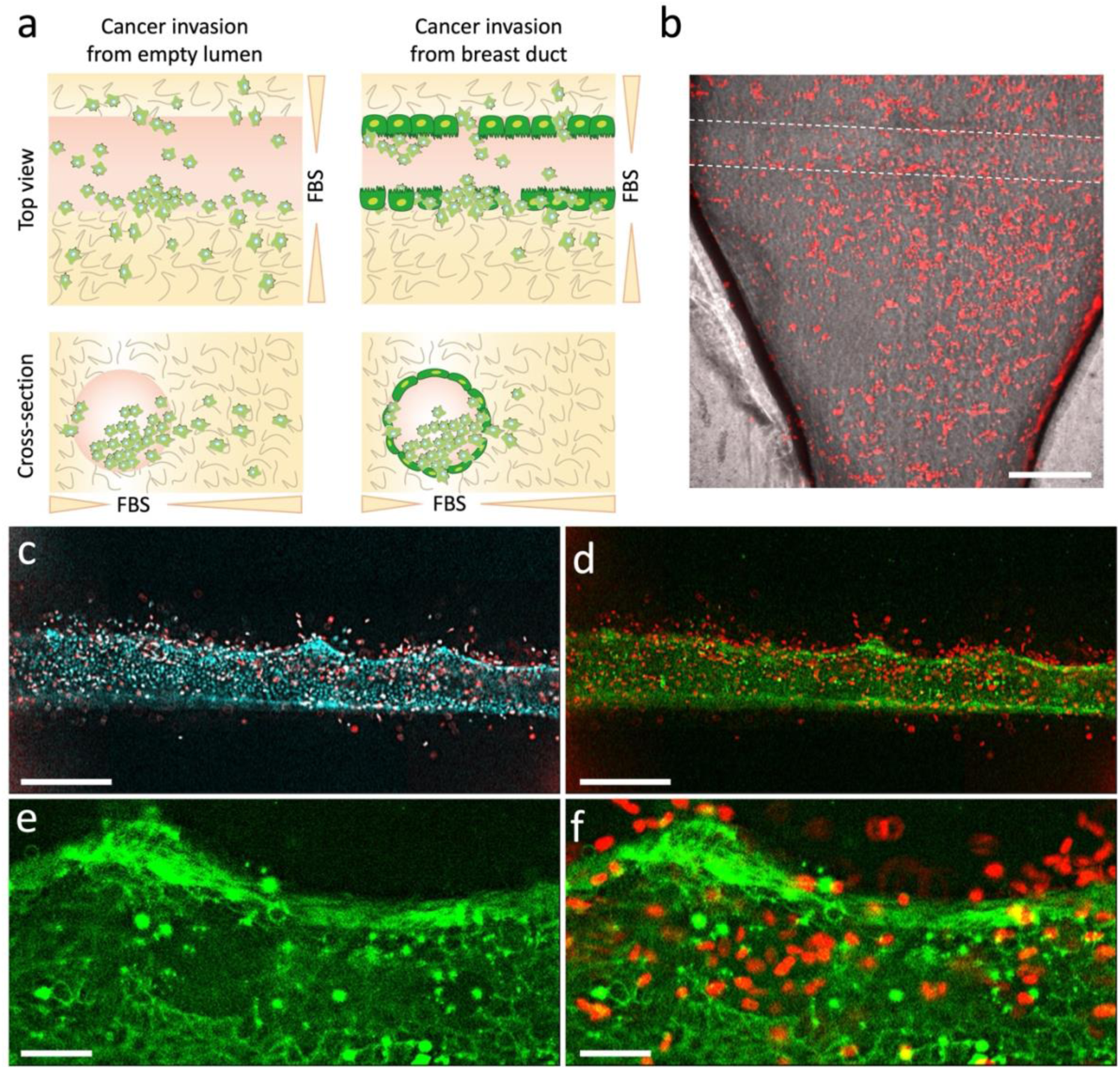
Breast DCIS and IDC on chip depending on initial cancer cell density. a) Schematic illustration of cancer cell invasion from (left) an empty channel versus (right) a breast duct lined with epithelial cells. The schematic shows the cancer invasion and the FBS source (as migratory trigger) from both top and cross-section view. The FBS was added to the media reservoirs (Figure 7.1a (ii)). b) MDA-MB-231 cancer cells (red) invading from an empty channel in absence of epithelial cells. Dashed lines show the edge of the channel within collagen gel. Scale bar: 200 μm. c, d) MDA-MB-231 cancer cell invasion from an epithelial duct lined with MCF10a cells. Inflorescent images stained for (c, d) MDA-MB-231 cancer cells in red, (c) cell nuclei in cyan, and (d) E-cadherin in green. The images are from a fixed sample after three days of invasion with a high initial cancer cell density. e, f) Magnified images from the edge of invasive breast ductal carcinoma in (d), stained for (e, f) E-cadherin in green, and (f) MDA-MB-231 cancer cells in red. Scale bars: 30 μm.

**Figure 3.**
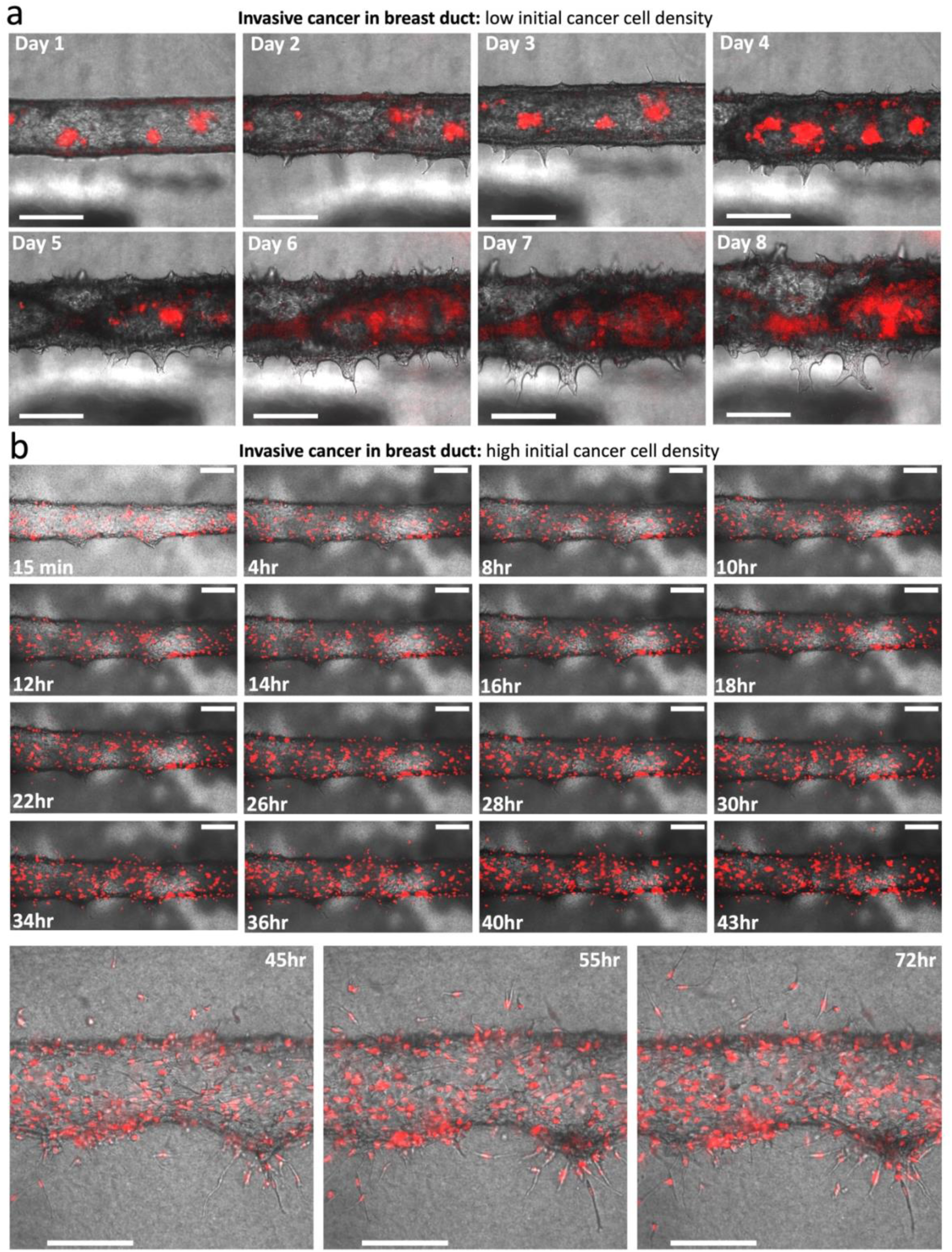
Invasive breast ductal carcinoma model depending on initial cancer cell density. a) Snapshots from a time-lapse recording of epithelial protrusions from a breast duct. The initial number of MDA-MB-231 cells was low, 2 · 10⁵ cells/ml. MDA-MB-231 cells are shown in red. Scale bars: 200 μm. See Supplementary video 2. b) Snapshots from a time-lapse recording of single MDA-MB-231 cancer cells invasion from a breast duct. See Supplementary video 3. The initial number of MDA-MB-231 cells was higher, 2 · 10⁶ cells/ml. The last three images show magnified images of cancer cells invading from the breast duct. MDA-MB-231 cells are shown in red. Scale bars: 100 μm.

**Figure 4.**
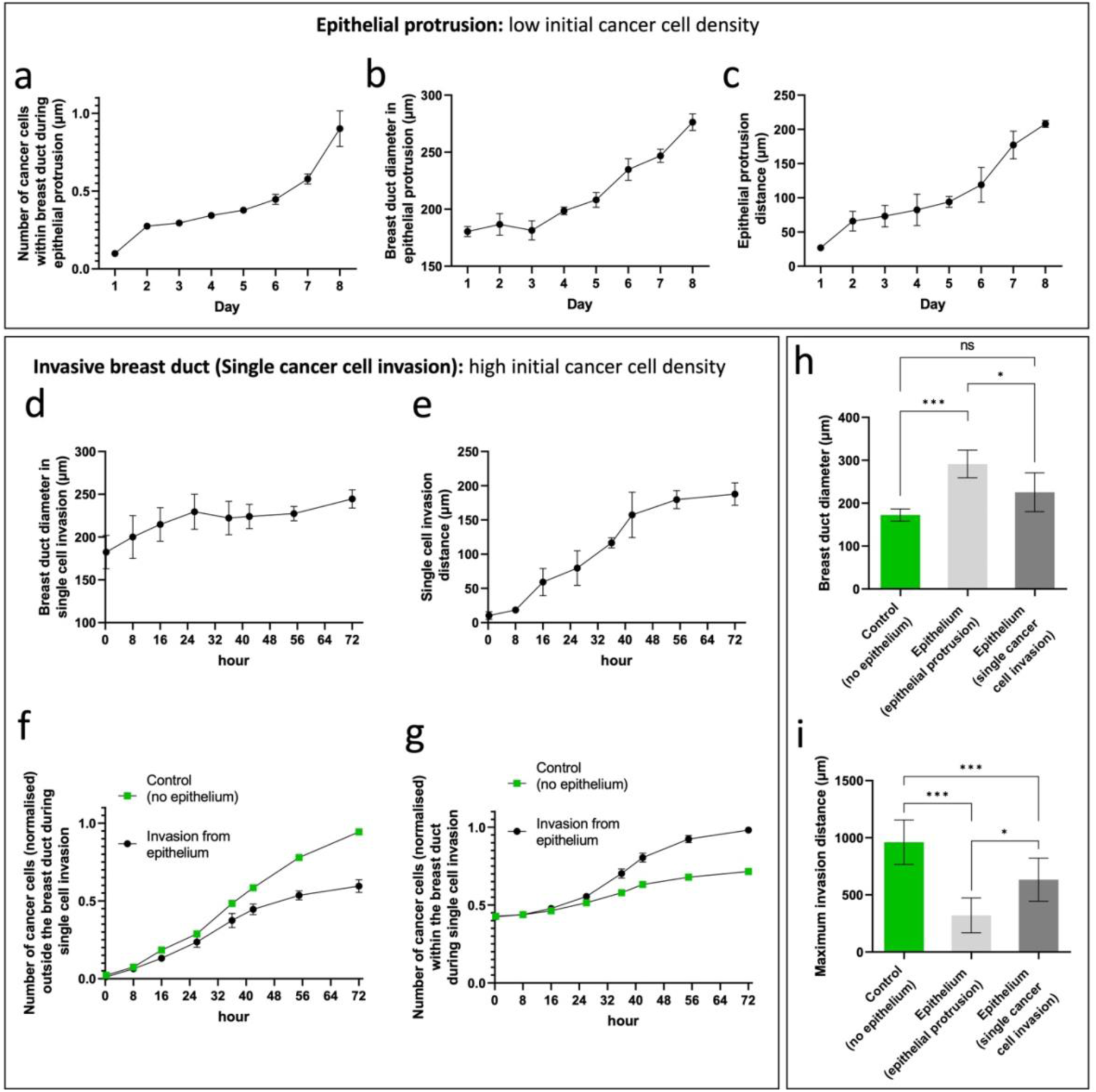
The regime of breast ductal carcinoma depends on the initial number of cancer cells within the epithelial duct. a–c) Epithelial protrusion with low initial cancer cell density. (a) Number of MDA-MB-231 cancer cells within the breast duct, (b) breast duct diameter, and (c) epithelial protrusion distance during 8 days of epithelial protrusion. The number of cancer cells was normalized by dividing by the highest value in the recorded data. d–g) Single cancer cell invasion with high initial cancer cell density. (d) Breast duct diameter, and (e) single cancer cell invasion distance during 72 hours of invasion. (f, g) Number of MDA-MB-231 cancer cells (f) outside and (g) within the breast duct during 72 hours of invasion. The number of cancer cells was normalized by dividing by the highest value in the recorded data. h, i) Breast duct diameter and maximum invasion distance in the control (in absence of the epithelial cells, and high initial number of cancer cells), epithelial protrusion (low initial cancer cell density), and single cell invasion (high initial cancer cell density) conditions within 8 days of invasion. *P < 0.05, ***P < 0.001, ****P < 0.0001 indicate statistical significance. ns indicates non-significance. One-way ANOVA, followed by Tukey post-hoc analysis were done to determine statistical significance.

At higher MDA-MB-231 seeding densities, a distinct invasion pattern was observed, characterized by single-cell dissemination through the epithelial barrier (Figure 3b, Supplementary video 3). In this regime, termed “single-cell invasion,” cancer cells populated the duct periphery and rapidly breached the epithelial lining. Despite ongoing invasion, the duct diameter remained largely unchanged during the three-day culture period (Figure 4d). Cancer cells migrated deeper into the surrounding matrix in this condition compared to the epithelial duct condition (Figure 4e). Furthermore, the number of non-invading cancer cells within the duct increased less in the control than in the epithelialized condition over 72 hours (Figure 4g), reinforcing the role of the epithelial barrier in modulating cancer cell behavior.

To further elucidate these invasion dynamics, we compared duct morphologies and invasion outcomes under three conditions: (1) epithelial protrusion (low cancer cell density), (2) single-cell invasion (high cancer cell density), and (3) control (no epithelium, high cancer cell density) across an 8-day culture period. The duct diameter remained stable in the control, but increased in the other two conditions, with a more pronounced expansion observed during epithelial protrusion (Figure 4h). The maximum cancer cell invasion distance was greatest in the control condition and was reduced in the presence of an epithelial barrier (Figure 4i). Among the two epithelialized conditions, invasion distance was higher in the single-cell regime than in the epithelial protrusion regime, indicating that collective epithelial migration progresses more slowly than single-cell dissemination, likely due to the maintenance of cell-cell junctions during group movement.

### Testing chemotherapies on invasive breast ductal carcinoma on chip

We employed the breast duct-on-chip platform to investigate the impact of chemotherapeutic agents on cancer cell invasion in breast ductal carcinoma. Specifically, we evaluated the effects of two widely used chemotherapies, Doxorubicin and Paclitaxel, on the invasion behavior of MDA-MB-231 cells (Figure 5a, b). Paclitaxel, a taxane-based chemotherapy agent, disrupts microtubule dynamics and halts cell division, thereby inhibiting the proliferation of cancer and other rapidly dividing cells [44][48]. It is commonly used as a first-line treatment in advanced breast cancer [44]. Despite its widespread clinical application, questions remain about optimal dosing strategies and pharmacokinetics. Paclitaxel is administered in various dosages and schedules depending on the clinical context, and its plasma concentration fluctuates over time due to hepatic clearance [44]. Additionally, Paclitaxel accumulates intracellularly within cancer cells [45,46], often reaching concentrations significantly higher than those observed in plasma, yet no established model exists to predict this intratumoral concentration from blood levels [44]. Biopsies are typically required for such measurements, which are not always feasible during treatment. Compounding the complexity, resistance to Paclitaxel frequently develops, reducing its therapeutic efficacy [47]. Doxorubicin, another commonly used chemotherapeutic agent in breast cancer, intercalates into DNA and inhibits topoisomerase II, thereby preventing DNA replication and transcription [48–50]. While effective in many cases, Doxorubicin is also associated with the development of drug resistance and may paradoxically enhance metastasis by upregulating pro-migratory factors in cancer cells [51,52]. Furthermore, the timing and dose-dependence of these effects remain poorly understood, especially in the context of combination regimens such as Doxorubicin with Paclitaxel [53,54].

**Figure 5.**
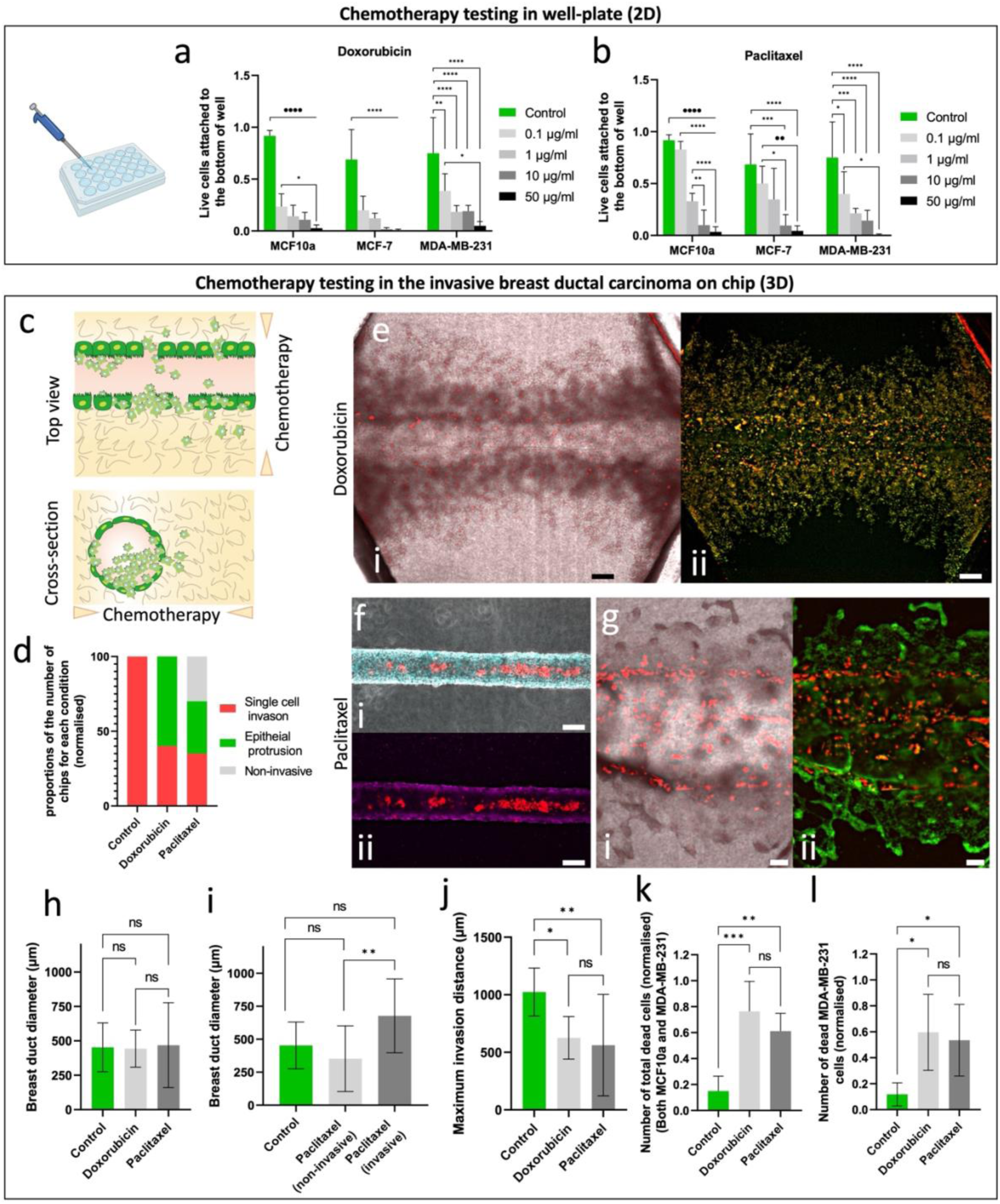
Testing chemotherapies on invasive breast ductal carcinoma on chip. a, b) Number of live cells (normalized) in response to various concentrations of (a) Doxorubicin and (b) Paclitaxel. These measurements were done in a well-plate for normal breast epithelial cells (MCF10a), non-invasive (MCF-7), and invasive (MDA-MB-231) breast cancer cells. c) Schematic illustration of testing chemotherapy in an invasive ductal carcinoma. d) Percentage of tests with specific regimes in response to 0.1 μg/ml of Doxorubicin and Paclitaxel. Here, we tested only invasive cancer cells (MDA-MB-231) cells in the breast duct lined with MCF10a cells. The initial cancer cell density was high (2 · 10⁶ cells/ml) in all drug testing experiments. e) Images of a non-invasive breast DCIS response to 0.1 μg/ml Paclitaxel. This is an example of a test in which cancer cells did not invade when treated with Paclitaxel. i = Overlapped image of phase contrast and fluorescent images (stained for cell nuclei in cyan and MDA-MB-231 cancer cells in red). ii = Fluorescent image stained for E-cadherin in magenta and cancer cells in red. Scale bars: 100 μm. f) Images of a model showing epithelial protrusion in response to 0.1 μg/ml Paclitaxel. i = Overlapped image of phase contrast and fluorescent images (stained for MDA-MB-231 cancer cells in red). ii = Fluorescent image stained for F-actin in green and cancer cells in red. Scale bars: 100 μm. g) Images of an IDC in response to 0.1 μg/ml Doxorubicin. i = Overlapped image of phase contrast and fluorescent images (stained for MDA-MB-231 cancer cells in red). ii = Fluorescent image stained for cell nuclei in green and cancer cells in red. Scale bars: 100 μm. h) Breast duct diameter in control, in comparison with application of 0.1 μg/ml Doxorubicin and 0.1 μg/ml Paclitaxel conditions. i) Breast duct diameter in control, in comparison with invasive and non-invasive samples treated with 0.1 μg/ml Paclitaxel conditions. j) Maximum invasion distance of MDA-MB-231 cancer cells in control, in comparison with application of 0.1 μg/ml Doxorubicin and 0.1 μg/ml Paclitaxel conditions. k, l) Number of (k) total dead cells (normalized) and (l) MDA-MB-231 dead cells (normalized) in control, in comparison with application of 0.1 μg/ml Doxorubicin and 0.1 μg/ml Paclitaxel conditions. *P < 0.05, **P < 0.01, and ***P < 0.001 indicate statistical significance. ns indicates non-significance. One-way ANOVA, followed by Tukey post-hoc analysis were done to determine statistical significance.

There is a critical need for advanced *in vitro* models that enable controlled testing of such therapies under physiologically relevant conditions. OoC systems offer this capability, allowing for long-term monitoring and spatially controlled drug delivery in an *in vivo*-like microenvironment. In this study, we first tested a range of Doxorubicin and Paclitaxel concentrations on three cell types in 2D culture. Both drugs demonstrated concentration-dependent cytotoxicity across all cell types (Figure 5a, b). Based on these findings, we selected MDA-MB-231 cells and a concentration of 0.1 μg/mL for both drugs for subsequent testing in the duct-on-chip system. Chemotherapeutics were introduced through the media reservoirs (Figure 1a(ii)) to generate a concentration gradient toward the breast duct (Figure 5c). In untreated control conditions, MDA-MB-231 cells exhibited characteristic single-cell invasion into the matrix (Figure 5d). However, drug treatment altered this invasion profile (Figure 5d–g). Notably, Doxorubicin treatment led to epithelial protrusion in 60% of samples and single-cell invasion in the remaining 40% (Figure 5e). Paclitaxel treatment resulted in complete inhibition of invasion in 30% of cases, while the remaining samples showed either collective epithelial protrusion (35%) or single-cell invasion (35%) (Figure 5f). In conditions with epithelial protrusion under Paclitaxel treatment, MDA-MB-231 cells were frequently observed aligning along the base of protrusions extending from the duct (Figure 5g). The average breast duct diameter remained unchanged in drug-treated conditions compared to the untreated control (Figure 5h). However, under Paclitaxel treatment, ducts with invasive carcinoma exhibited significantly larger diameters than those without invasion (Figure 5i). Maximum invasion and protrusion distances were reduced under both drug treatments compared to controls (Figure 5j). Moreover, the number of dead cells, both total and cancer-specific, increased significantly following drug administration (Figure 5k, l). This was consistent with our well-plate experiments, where all cell types, including normal epithelial and cancer cells, showed similar sensitivity to increasing drug concentrations. This outcome was expected, as both Paclitaxel and Doxorubicin target dividing cells, regardless of their malignant status.

This cytotoxicity may have contributed to certain phenotypes observed in the duct-on-chip system. For instance, the observed increase in dead cells among both epithelial and cancer cell populations suggests a general inhibition of proliferation. In future studies, using patient-derived tumor cells could improve clinical relevance. Additionally, manipulating epithelial cell proliferation through genetic modification or targeted inhibitors may help decouple epithelial contributions from cancer-driven invasion. Such interventions could clarify the extent to which ductal geometry and epithelial behavior influence invasion dynamics and therapeutic response. Finally, ductal expansion observed in this model, approximately 2-to 3-fold under control conditions, may reflect *in vivo* processes, such as tumor-induced local inflammation or hyperplasia. While neither drug treatment further increased duct diameter (Figure 5h), the tumor microenvironment (TME) *in vivo* is more complex and includes immune and stromal cells that influence cancer progression. Incorporating immune cells into the extracellular matrix or introducing inflammatory cytokines into the culture media may further improve this model’s fidelity and provide insight into the interplay between inflammation, invasion, and therapeutic response.

### Intraductal migration of cancer cells in non-invasive ductal carcinoma-on-chip

Given the occurrence of non-invasive ductal carcinoma in both low cancer cell density conditions (Figure 3a) and in certain Paclitaxel-treated cases (Figure 5d, e), we further investigated the behavior of MDA-MB-231 cancer cells within the breast duct under these conditions. Remarkably, we observed a flow-like translational motion of cancer cell clusters along the ductal axis (Figure 6). This movement appeared to be guided by collective epithelial migration, with cancer clusters passively sliding along the migrating epithelial monolayer (Figure 6a and Supplementary video 4). In one representative experiment, cancer clusters moved unidirectionally from left to right, while the epithelial cell motion reversed direction approximately 15.5 hours into the observation (Figure 6a). A similar pattern was seen in Paclitaxel-treated non-invasive samples, where cancer clusters migrated leftward before reversing direction at the 12-hour mark (Figure 6b and Supplementary video 5). These findings suggest dynamic, coordinated behavior between epithelial and cancer cell populations during the non-invasive phase of ductal carcinoma. The ductal epithelium appeared to act as a physical barrier, preventing lateral invasion into the surrounding extracellular matrix and potentially limiting drug diffusion to the tumor clusters. Moreover, the epithelial flow may mimic an *in vivo* mechanism of early-stage intraductal carcinoma, such as ductal carcinoma *in situ* (DCIS), where malignant cells proliferate within the ductal lumen without penetrating the BM. Understanding these dynamics is essential to elucidate the transition mechanisms from non-invasive to invasive stages, such as the progression from DCIS to IDC. In our model, cancer cells primarily remained clustered at the ductal center, although single cell shedding at the ductal edges was also observed (Figure 6c). This shedding may represent an early event in lateral invasion, where individual cells adhere to the epithelial BM and potentially breach it. In one notable instance, we observed a cancer cell exiting and then re-entering the ductal epithelium (Figure 6c, yellow arrows). Such behavior reflects processes in DCIS, where cells can detach into the lumen, contributing to tumor burden and, potentially, later escape into surrounding tissue. Previous studies suggest that these shed cells may re-adhere to the BM or exploit weakened regions to initiate invasion, a mechanism potentially contributing to the onset of invasive disease.

**Figure 6.**
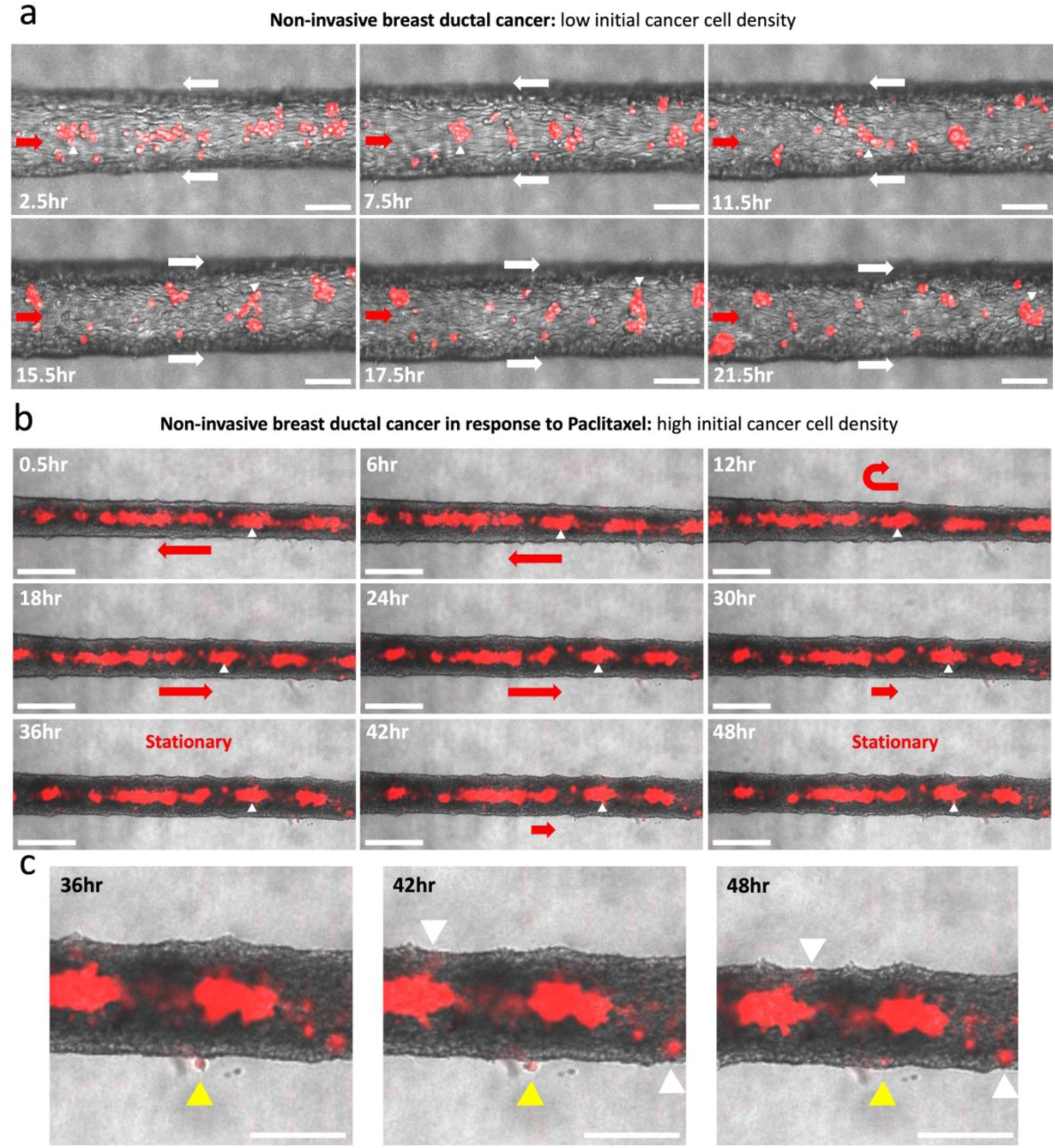
Intraductal migration of cancer cells in non-invasive breast ductal carcinoma. a) Snapshots from time-lapse recordings of non-invasive ductal carcinoma with low initial cancer cell density. The initial number of MDA-MB-231 cells was 2 · 10⁵ cells/ml. MDA-MB-231 cells are shown in red. Red arrows show the movement direction of the MDA-MB-231 cancer cell clusters, white arrows, the movement of the epithelial cells (MCF10a) at the edges. White triangle markers point at the same cluster in every snapshot, showing its intraductal movement. Scale bars: 100 μm. See Supplementary video 4. b) Snapshots from time-lapse recordings of non-invasive breast ductal carcinoma treated with 0.1 μg/ml Paclitaxel. MDA-MB-231 cells are shown in red. The initial number of MDA-MB-231 cells was 2 · 10⁶ cells/ml. MDA-MB-231 cells are shown in red. Red arrows show the movement direction of the MDA-MB-231 cancer cell clusters. White triangle markers point at the same cluster in every snapshot, showing its intraductal movement. See Supplementary video 5. c) Zoomed images from the last three snapshots in (b) show cancer cells within the breast duct shedding into the epithelium at the edges. Yellow arrows indicate a single cancer cell shedding out of the breast duct and then returning. Scale bars: (a, b) 200 μm, and (c) 100 μm.

Our duct-on-chip model offers a powerful platform for investigating these dynamics, particularly with the incorporation of more clinically relevant cell types. The uncontrolled proliferation of immortalized cell lines, such as MCF10A, may influence epithelial flow and cancer cell behavior. Utilizing primary patient-derived epithelial and cancer cells or cell lines with inducible proliferation control could improve model fidelity and better reflect patient-specific biology. While our current system successfully mimics key features of both DCIS and IDC, it also presents limitations that should be addressed in future research. The model is currently incompatible with softer or low concentration hydrogels due to the need for stable structures during the needle-pulling step. Future iterations could incorporate structural modifications or alternative biomaterials to support softer matrix environments. Additionally, automating the needle-removal step may minimize variability introduced by manual handling. Another limitation of our study is the exclusive use of established cell lines, which may not fully capture the heterogeneity and microenvironmental complexity of clinical tumors. Incorporating immune cells, stromal components (e.g., cancer-associated fibroblasts), and inflammatory factors could expand the model’s utility in studying tumor microenvironment interactions [2,55]. Furthermore, although we tested chemotherapies at a single concentration, a broader concentration-dependent analysis would provide deeper insights into dose-response relationships. Our treatment regimen (three days of drug exposure over an eight-day culture period) may not fully replicate clinical protocols. Future studies could integrate patient-derived materials and personalized treatment schedules to improve translational relevance. Ultimately, early integration of chemotherapy testing into chip development is crucial for advancing these models toward functional implementation in high-throughput drug screening platforms. We believe our ductal carcinoma-on-chip model lays a foundational step toward that goal.

## Conclusion

The inclusion of an epithelial layer in models simulating ductal carcinoma and invasion is essential for elucidating the transition from ductal carcinoma *in situ* (DCIS) to invasive breast cancer. In this study, we introduced a versatile yet accessible ductal carcinoma-on-chip model that enables detailed investigation of invasion dynamics and therapeutic responses. We found that the mode of invasion, whether through epithelial protrusions or single-cell dissemination, depended heavily on the initial cancer cell density. Additionally, both the invasion distance and ductal morphology varied across conditions, highlighting the importance of cellular composition and cell-type ratios in determining experimental outcomes. Using this model, we tested two frontline breast cancer chemotherapies, Doxorubicin and Paclitaxel. Both drugs significantly altered the invasion patterns and duct architecture, with Paclitaxel notably halting invasion and protrusion in 30% of tested conditions. However, these therapies also affected all proliferative cell types within the system, not just cancer cells, underlining a limitation common to many current treatments. Our platform offers a powerful system for studying early-stage ductal carcinoma progression and holds potential for evaluating diverse therapeutic agents and biological factors involved in cancer development. These advanced platforms provide valuable, scalable tools for investigating cancer biology and accelerating the development of next-generation therapies, including those targeting complex features of solid tumors.

## Supporting information

Supplementary video 1. Breast duct-on-chip model (Time lapse). Normal breast epithelial cells (MCF10a) lining a tubular channel within collagen I.

Supplementary video 2. Invasive breast ductal carcinoma (Time lapse). Breast cancer cells (MDA-MB-231, red) within a breast duct-on-chip model lined w

Supplementary video 3. Invasive breast ductal carcinoma (Time lapse). Breast cancer cells (MDA-MB-231, red) invading from a breast duct-on-chip model

Supplementary video 4. Non-invasive breast duct on chip model (Time lapse). Breast cancer cells (MDA-MB-231, red) moving inside a breast duct-on-chip

Supplementary video 5. Non-invasive breast duct on chip model (Time lapse) in response to Paclitaxel. Breast cancer cells (MDA-MB-231, red) moving ins

## Acknowledgements

This work was supported by the European project Moore4Medical [10028031], and by the Dutch Research Council NWO (grant number Science-XL 2019.022, ‘The Active Matter Physics of Collective Metastasis’). Moore4Medical (https://moore4medical.eu/) has received funding within the Electronic Components and Systems for European Leadership Joint Undertaking (ECSEL JU) in collaboration with the European Union’s H2020 Framework Programme (H2020/2014-2020) and National Authorities, under grant agreement H2020-ECSEL-2019-IA-876190.

## Author contributions

Conceptualization: M.J.; methodology: M.J.; validation: M.J.; investigation: M.J.; resources: J.M.J.d.T.; writing – original draft: M.J.; writing – review & editing: M.J., and J.M.J.d.T.; visualization: M.J.; supervision: J.M.J.d.T.; project administration: J.M.J.d.T.; funding acquisition: J.M.J.d.T. All authors have read and agreed to the published version of the manuscript.

## Declaration of interests

The authors declare no competing interests.

## Methods

### Fabrication of microfluidic chip

The mold design was made in Siemens NX (Siemens AG) and then transferred to PreForm software (Formlabs). A durable resin cartridge was inserted into a Low Force Stereolithography 3D printer (both from Formlabs), and the printing was initiated. When the print was complete, the platform was placed in Form Wash (Formlabs) for 30 minutes to wash the uncured resin in isopropyl alcohol, and the resin was then fully cured in Form Cure (Formlabs) for 1 hour. The chip was made from a PDMS layer that was bonded to a glass slide using the following fabrication process. First, PDMS (Sylgard® 184 base with curing agent, both from Merck, at a 10:1 w/w ratio) was cast on the resin mold, degassed for 15 minutes, and cured at 65 °C. The poured PDMS weight was measured and dosed upfront to obtain a final layer thickness of 5 mm. After curing, the PDMS slabs were gently peeled off, and the edges were trimmed using a cutter. To create inlet and outlet access holes, 5 mm biopsy punchers (KAI biopsy punch, 4560146922619) were used. The channels were then sealed by bonding the PDMS layer to a 25 mm × 75 mm glass slide (VWR). For this, the PDMS slab (features facing up) and the glass slide were both exposed to 20 W air plasma for 30 seconds using a plasma asher (Emitech, K1050X). To achieve full bonding, the assembled complex underwent a thermal treatment at 65 °C for a minimum of 1 hour.

### Chip preparation and loading, and culture

The chips were autoclaved for sterility and all the next steps were conducted in a safety cabinet. The chips were located in a Petri dish (100 mm) for incubation, and a smaller Petri dish (35 mm) filled with Phosphate buffered solution (PBS) was placed next to the chip to increase the environmental humidity and to avoid excessive evaporation. The chamber of the chip was filled with 0.05% w/v dopamine hydrochloride (poly-dopamine; Thermo Scientific Chemicals via Fisher Scientific) for 1 hour at RT to enhance the hydrogel to PDMS attachment later. The polydopamine solution was aspirated out and the chamber was washed with PBS two times. All PBS was aspirated out, making sure that the chamber was empty. Steel acupuncture needles (160 μm diameter, Seirin) were introduced into the device through the lumen inlet of the chamber. The needle easily penetrated the PDMS to reach the lumen reservoir, and after needle extraction, the PDMS recovered with no leakage issues. Collagen type I (rat tail, 10 mg/mL, ibidi GmbH, Germany) solution was buffered with 10× PBS and titrated to a neutral pH with 0.1 M NaOH, and brought to a final concentration of 5 mg/mL collagen I. 100 μL of collagen-cell solution was transferred to the chamber of the microfluidic device and polymerized for 45 min at 37 °C, 5% CO₂. After polymerization, needles were gently extracted to create a hollow lumen. 10 μL of Matrigel solution (1:100 v:v in PBS) was gently injected in the channel and sample was placed in incubator to coat the channel with BM proteins. Then, the Matrigel solution was aspirated out. Next, 10 μL of MCF10a cells (4 · 10⁶ cells/mL) were pipetted into the lumens and incubated for 45 minutes at 37 °C, 5% CO₂ to facilitate attachment to the luminal base. After incubation, the reservoirs of the lumen were filled with MCF10a medium to wash out the non-attached cells. The media from these reservoirs were removed and 10 μL of PBS was pipetted into the lumen. Lateral hydrogel reservoirs were filled with MCF10a medium. After 24 hours of incubation, the MDA-MB-231 cells (low density 2 · 10⁵ or high density 2 · 10⁶ cells/mL) were added within the breast duct. The media composed of a 1:1 mix of MCF10a and MDA-MB-231 culture media, supplemented with 20% FBS, was added to the lateral gel reservoirs. Chips were cultured for 8 days at 37 °C, 5% CO₂. Media was exchanged daily through the gel reservoirs.

### Chemotherapy preparation and testing in 2D

Doxorubicin-Hydrochloride (Merck, D1515-10MG) was purchased in powder, and then dissolved in Dimethyl Sulfoxide (DMSO) to prepare the stock solution (10 mg/mL). The solution was further diluted in relevant cell culture media to prepare the final concentration. The DMSO concentration reduced to less than 0.1% in all concentrations. Paclitaxel (Merck, Y0000698, 40 mg) was purchased in powder, and then dissolved in Dimethyl Sulfoxide (DMSO) to prepare the stock solution (40 mg/mL). The solution was further diluted in relevant cell culture media to prepare the final concentration. The DMSO concentration reduced to less than 0.1% in all concentrations. The MCF10a, MCF-7 and MDA-MB-231 cells (1·10⁵ cells/mL) were cultured in 24 well-plates (Corning). Next day, various concentrations (0.1, 1, 10 and 50 μg/mL) of Doxorubicin and Paclitaxel diluted in relevant culture media were added to the plates and incubated for 72 h. Next, the cell nuclei were stained for live/dead using the ReadyProbe cell viability kit (Thermo Fisher, R37609). Since most of the dead cells were detached from the bottom of the wells and were suspended in media, we only imaged the bottom of the well plates. The number of live cells at the bottom of the wells were plotted, as shown in Figure 5a, b. The results are recorded from two experimental replicas, and 5 regions of interest per condition per replica.

### Testing chemotherapy drugs in Breast ductal carcinoma-on-chip

When MDA-MB-231 cells were seeded into the breast duct channel, the media (and drugs) were administered through lateral gel reservoirs. A drug concentration of 0.1 μg/mL was prepared in a 1:1 mixture of MCF10a and MDA-MB-231 culture media, supplemented with 20% FBS. The media was refreshed daily. Drug-supplemented media was introduced into the chips on the first, third, and sixth days of the culture period, while drug-free media was used for the remaining days of the 8-day culture period. On day 8, the samples were fixed and prepared for staining and imaging.

### Immunostaining and live imaging

The cell samples were fixed using 1.85% v/v formaldehyde (Merck, 1040031000) in PBS (Westburg, LO BE02-017F) for 15 minutes. The samples were permeabilized by exposing them to 0.5% v/v Triton X-100 (Merck, 108603) solution in PBS for 15 minutes. Samples were blocked via exposure to blocking solution (1% w/v Bovine Serum Albumin) (BSA) (Sigma-Aldrich, 9048-46-8), and incubated at room temperature for 1 hour. The cell nuclei were stained using NucBlue Fixed Cell ReadyProbe Reagent (Thermo Fisher, R37606). BM staining: Anti-collagen IV polyclonal (PA1-28534, 5 μg/ml), Anti-laminin monoclonal (11578772, 10 μg/ml) were from Invitrogen. Goat anti-rat IgG (ab150167, 2 μg/ml) was purchased from Abcam and Donkey anti-rabbit IgG (R37118, 2 drops/ml of PBS) secondary antibody was purchased from Thermo Fisher. Invasion assays and chemotherapy staining: F-actin using ActinGreen 488 ReadyProbes Reagent (Thermo Fisher, R37110). Mouse anti-E-Cadherin (Fisher Scientific, 10383223) and goat anti-Mouse (Thermo Fisher Scientific, A28180). The time lapse images and videos were recorded using inside incubator microscopes (Axion Biosystems-Lux3). Fixed samples were imaged by a Thunder-imaging system (Leica Microsystems) fluorescent microscope.

### Data analysis and schematics

The breast duct diameter was measured using Fiji. Invasion distance was also measured in Fiji, with the baseline defined as the centerline of the duct. Cell counts were performed in Fiji, and some data were normalized by dividing the values by the maximum value in the dataset. For each experimental condition, at least three replicate samples were analyzed. The schematics were made in Adobe Illustrator and Siemens NX (Siemens AG) software. Some schematic items were taken from the BioRender library. GraphPad Prism was used for generating graphs. One-way ANOVA, followed by Tukey post-hoc analysis, was done to determine statistical significance. P values < 0.05 were considered statistically significant.

### Statistics

Statistical analysis was performed using two-way ANOVA with Tukey’s multiple comparisons test in GraphPad Prism 9 software (GraphPad Software Inc., San Diego, CA, USA). Differences were considered to be significant when p < 0.05.

## Supplementary

**Supplementary video 1.** Breast duct-on-chip model (Time lapse). Normal breast epithelial cells (MCF10a) lining a tubular channel within collagen I.

**Supplementary video 2.** Invasive breast ductal carcinoma (Time lapse). Breast cancer cells (MDA-MB-231, red) within a breast duct-on-chip model lined with normal epithelial cells (MCF10a). At low initial cancer cell density within the duct, these cells did not invade from epithelium but induced collective epithelial protrusion.

**Supplementary video 3.** Invasive breast ductal carcinoma (Time lapse). Breast cancer cells (MDA-MB-231, red) invading from a breast duct-on-chip model lined with normal epithelial cells (MCF10a). At high initial cancer cell density within the duct, these cells invaded through epithelium into the collagen I gel (single cell invasion).

**Supplementary video 4.** Non-invasive breast duct on chip model (Time lapse). Breast cancer cells (MDA-MB-231, red) moving inside a breast duct-on-chip model lined with normal epithelial cells (MCF10a).

**Supplementary video 5.** Non-invasive breast duct on chip model (Time lapse) in response to Paclitaxel. Breast cancer cells (MDA-MB-231, red) moving inside a breast duct-on-chip model lined with normal epithelial cells (MCF10a). The cancer cell clusters move inside the duct but do not invade out.

## Notes

### Competing Interest Statement

The authors have declared no competing interest.

